# A phased, chromosome-scale genome of ‘Honeycrisp’ apple (*Malus domestica*)

**DOI:** 10.1101/2022.08.24.505160

**Authors:** Awais Khan, Sarah B. Carey, Alicia Serrano, Huiting Zhang, Heidi Hargarten, Haley Hale, Alex Harkess, Loren Honaas

## Abstract

‘Honeycrisp’ is one of the most valuable apple cultivars grown in the United States and a popular breeding parent due to its superior fruit quality traits, high levels of cold hardiness, and disease resistance. However, it suffers from a number of physiological disorders and is susceptible to production and postharvest issues. Although several apple genomes have been sequenced in the last decade, there is still a substantial knowledge gap in understanding the genetic mechanisms underlying cultivar-specific traits. Here we present a fully phased, chromosome-level genome of ‘Honeycrisp’ apples, using PacBio HiFi, Omni-C, and Illumina sequencing platforms. Our genome assembly is by far the most contiguous among all the apple genomes. The sizes of the two assembled haplomes are 674 Mb and 660 Mb, with contig N50s of 32.8 Mb and 31.6 Mb, respectively. In total, 47,563 and 48,655 protein coding genes were annotated from each haplome, capturing 96.8-97.4% complete BUSCOs in the eudicot database, the most complete among all *Malus* annotations. A gene family analysis using seven *Malus* genomes shows that a vast majority of ‘Honeycrisp’ genes are assigned into orthogroups shared with other genomes, but it also reveals 121 ‘Honeycrisp’-specific orthogroups. We provide a valuable resource for understanding the genetic basis of horticulturally important traits in apples and other related tree fruit species, including at-harvest and postharvest fruit quality, abiotic stress tolerance, and disease resistance, all of which can enhance breeding efforts in Rosaceae.

## Main Content

### Background

Apples are the most consumed fruit in the United States (www.ers.usda.gov). The annual estimated total value of the US apple industry is $21 billion, with five cultivars alone accounting for 2/3 of production (in order of proportion): ‘Gala’, ‘Red Delicious’, ‘Honeycrisp’, ‘Granny Smith’, and ‘Fuji’ (www.usapple.org). Of these, ‘Honeycrisp’ is by far the most valuable - it has roughly twice the value per pound of the next most valuable cultivar, ‘Fuji’ [1]. ‘Honeycrisp’ is appreciated by consumers, and therefore by the US apple industry, for its superior flavor and crisp juicy texture. Importantly, properly stored ‘Honeycrisp’ fruit is well-preserved for several months [2,3]. Additionally, this cultivar shows high levels of cold hardiness [4] and resistance to apple scab, the most economically important fungal disease of apples worldwide [5]. ‘Honeycrisp’ was bred at the University of Minnesota in the 1960s aiming to obtain cold hardy cultivars with high-quality fruit; it was released in 1991 [6] (Figure 1A). Recent genome-wide analysis (following the resolution of the ‘Honeycrisp’ pedigree [7,8]) showed that the genetic background of ‘Honeycrisp’ is distinct from other important apple cultivars in the US. This is highlighted by the success of ‘Honeycrisp’ as a source of interesting genetic diversity in apple breeding programs worldwide to enhance texture, storability, and improved disease resistance [3,5,7,9,10]. In fact, nine new cultivars derived from ‘Honeycrisp’ are already on the market.

**Figure 1:**
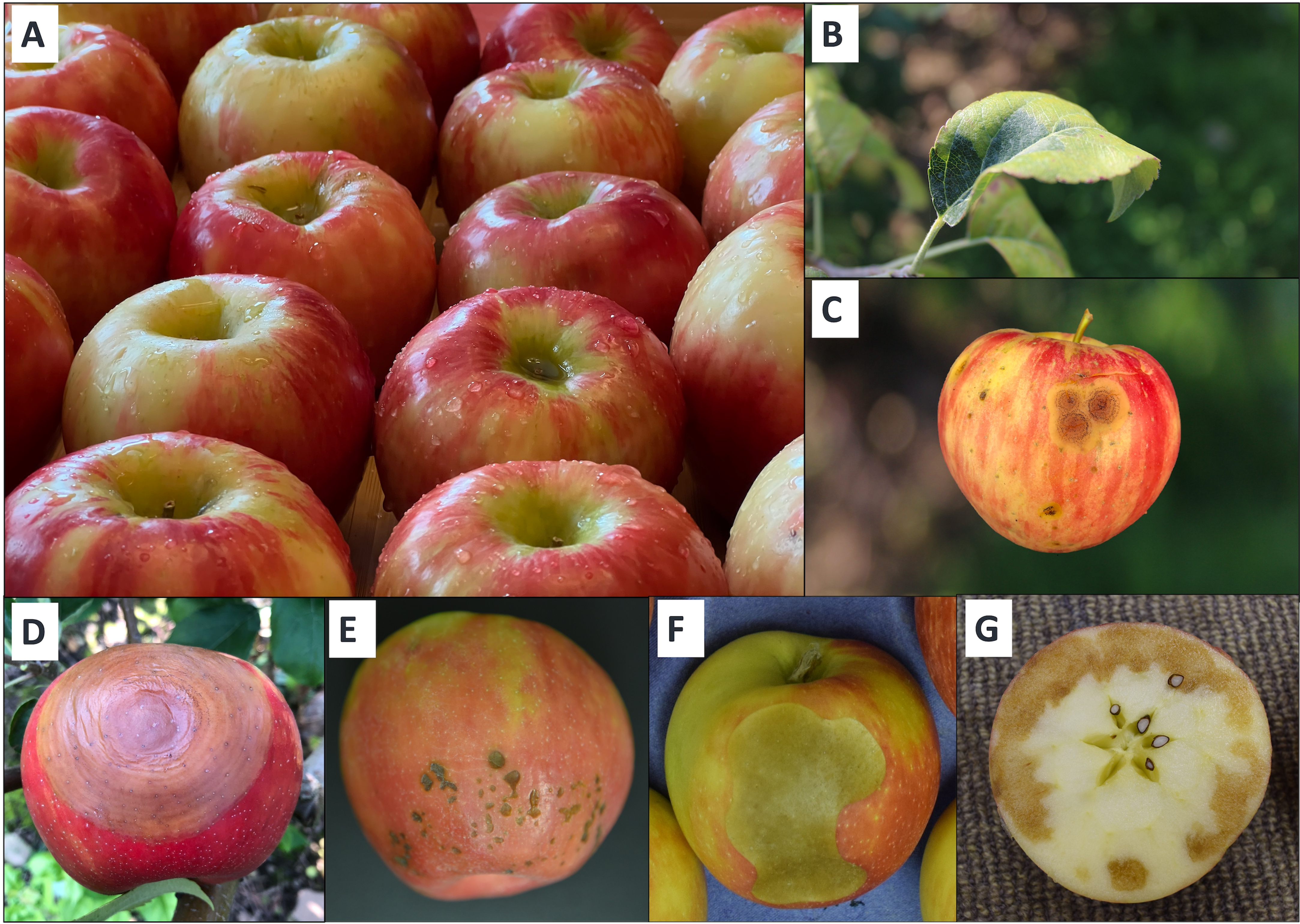
‘Honeycrisp’ is a highly desirable apple cultivar, appreciated by consumers for its superior flavor, texture, and visual appeal (A). To fulfill high-market demand for ‘Honeycrisp’, growers need to optimize production traits, and address several diseases and physiological disorders of concern to ensure high levels of production and fruit quality, including zonal leaf chlorosis (B), fungal diseases like the bitter rot pathogen complex (*Colletotrichum gloeosporiodes* and *C. acutatum*) (C) the black rot pathogen (*Botryosphaeria obtuse*) (D), as well as postharvest storage disorders like bitter pit (E) soft scald (F), and soggy breakdown (G).

Disease resistance, critical for sustainable apple production, has historically been less important due to a market dominated by modern cultivars bred primarily for fruit quality and intensive conventional production systems [11]. Most apple cultivars grown commercially in the US are susceptible to fungal diseases such as apple scab. In temperate and humid regions around the world, frequent applications of fungicides are necessary, contributing significantly to production costs, and to negative human health and environmental impacts [12]. ‘Honeycrisp’ is resistant to apple scab and importantly, this cultivar’s ability to retain crispness and firmness during storage is one of the most outstanding traits of ‘Honeycrisp’ fruit [13]. However, there are other production issues with ‘Honeycrisp’ that present challenges for apple growers (Figure 1E-G). ‘Honeycrisp’ needs a carefully designed nutrient management program during the growing season for optimal production and fruit quality, especially to limit the occurrence of the physiological disorder bitter pit [3]. ‘Honeycrisp’ trees also have greater tendency to develop zonal leaf chlorosis, a physiological disorder that reduces photosynthetic capacity [14]. However, in the Pacific Northwest (PNW), where a large majority of the ‘Honeycrisp’ apples are grown in the US (www.nwhort.org) due in part to low disease pressure, postharvest issues during long-term storage pose substantial challenges to producers.

The total cullage of ‘Honeycrisp’ fruit is likely among the highest of apple cultivars due to its susceptibility to various postharvest physiological disorders, which have complex etiologies that are poorly understood, and include bitter pit, soft scald, soggy breakdown, and CO_2_ injury [15–18]. Postharvest technologies have been developed and deployed to mitigate these disorders [19–21]. However, the efficacy of postharvest treatments can be affected by many factors such as pre-harvest orchard management and at-harvest fruit maturity, a key factor in the maintenance of postharvest apple fruit quality. Growers must balance the acquisition of certain fruit quality characteristics (*e.g.* size, color, flesh texture, and sugar content), while attempting to minimize risk for maturity-linked losses in quality that may occur in the supply chain [22]. This balancing act for maximizing at-harvest fruit quality and long-term cold storage potential in controlled atmospheres is especially difficult for ‘Honeycrisp’.

To maximize both our understanding of genetic mechanisms driving important ‘Honeycrisp’ traits and to assist tree fruit breeders, high quality genomes are required [23]. Indeed, in the last decade since ‘Golden Delicious’ was sequenced [24], a large number of genes and QTLs linked to fruit disease resistance, quality traits, and abiotic stress tolerance in apples have been identified [5,25,26]. Recent high-quality genomes of ‘Gala’, the double haploid ‘Golden Delicious’, and the triploid ‘Hanfu’ provide genomic resources for apple genetics and breeding [27–29]. These studies have identified targeted genomic regions for the development of diagnostic molecular markers to breed disease resistant apple cultivars with good fruit quality [30]. However, the fact remains that traditional apple breeding is a resource-intensive and time-consuming process [9,26,30] and there are still substantial gaps in our knowledge of genetic mechanisms involved in many important apple traits. In this manuscript we report a phased, chromosome-level genome assembly of the ‘Honeycrisp’ apple cultivar generated from PacBio HiFi and Dovetail Omni-C, plus a high-quality annotation – thus providing one of the best genome resources available for apples to date.

## Methods

### PacBio HiFi sequencing

Cuttings of dormant wood were collected from ‘Honeycrisp’ trees growing in the experimental orchard at Cornell AgriTech (Geneva, NY, USA). The cuttings were placed in water in the greenhouse until leaves began emerging from the buds, and thereafter placed in the dark for two days. Young, dark-adapted leaves were collected and shipped on dry ice to the DNA Sequencing and Genotyping Center at the University of Delaware (DL, USA) for DNA extraction and Single Molecule Real Time (SMRT) PacBio (Pacific BioSciences) sequencing.

High-molecular-weight (HMW) genomic DNA was extracted using a DNeasy Plant Mini Kit (Qiagen) according to the manufacturer's protocol. HMW genomic DNA was sheared to 15 kb fragments, and the HiFi library was prepared using SMRTbell Express Template Prep Kit 2.0 and the DNA/Polymerase Binding Kit 2.0 (Pacific Biosciences) according to the manufacturer’s protocol. The sequencing library was size-selected using Sage Blue Pippin (Sage Sciences) to select fragment sizes of >10 kb to ensure removal of smaller fragments and adapter dimers. The library was sequenced on a PacBio Sequel II instrument in CCS/HiFi mode with two SMRT cells with 2 hours pre-extension and 30-hour movie times. Read length distribution and quality of all HiFi reads was assessed using Pauvre v0.1923 (https://github.com/conchoecia/pauvre).

To scaffold the genome using chromatin conformation sequencing, 1 g of flash-frozen young leaf material was harvested from ‘Honeycrisp’ trees at the Washington State University Sunrise Research Orchard near Rock Island, WA USA and shipped to the HudsonAlpha Institute for Biotechnology in Huntsville, AL USA. The sequencing library was prepared using the Dovetail Genomics Omni-C kit and was sequenced on an Illumina NovaSeq 6000 with PE150 reads. A subset of 1 million read pairs were used as input for Phase Genomics *hic_qc* to validate the overall quality of the library (https://github.com/phasegenomics/hic_qc).

### Phased haplome assembly and scaffolding

The expected genome size, heterozygosity, and percent of repeats was assessed by generating 21-mer sequences from the raw HiFi data with Jellyfish v2.3.0 [31] and GenomeScope2 [32,33]. HiFi reads were assembled into contigs using hifiasm v0.16.1 [34,35], with the Hi-C integration mode that incorporated Dovetail Omni-C reads for phasing. Both haplomes of the assembly were scaffolded into chromosomes using the Juicer pipeline v1.6 [36], where the Omni-C reads were mapped separately to both hifiasm haplomes [35,37] with the parameter “-s none”. The Omni-C data was subset to ~100x coverage and the 3D-DNA v201008 scaffolding pipeline [38] was run with options “--editor-saturation-centile 10 --editor-coarse-resolution 100000 --editor-coarse-region 400000 --editor-repeat-coverage 50”. Contact maps were manually edited using the Juicebox Assembly Tools (JBAT) v1.11.08 [36] to produce the expected 17 chromosomes per haplome. Contigs that contained assembled telomeres were correctly oriented to the terminal ends by searching for the TTTAGGG repeat (or the reverse complement CCCTAAA) using the analyze_genome function of GENESPACE [39]. The chromosomes were numbered and oriented using haplome A of the ‘Gala’ assembly [27]. Genome quality and completeness was assessed using benchmarking universal single-copy gene orthologs (BUSCO v5.2.2) [40] with the “eudicots_odb10” database. Haplome completeness was also assessed using Merqury v1.3 [41].

### Transcriptome sequencing

To facilitate gene annotation, total RNA was isolated from various tissues harvested from ‘Honeycrisp’, ‘Red Delicious’, and ‘Granny Smith’ apple trees grown at the Washington State University (WSU) Sunrise Research Orchard near Rock Island, WA USA, ‘Gala’ and ‘WA38’ apple trees grown at the WSU and USDA-ARS Columbia View Research Orchard near Orondo, WA USA, and ‘D’Anjou’ pear trees grown at the WSU Tree Fruit Research and Extension Center Research Orchard in Wenatchee, WA USA using a modified CTAB/Chloroform extraction [42]. Total RNA was assessed for quality (RNA integrity number (RIN) ≥8) and purity (A260/280 >1.8). Sources for all RNA are available in Table 3. 2 μg of total RNA was used to construct Illumina TruSeq stranded libraries following manufacturers’ instructions. Libraries were sequenced on an Illumina NovaSeq 6000 with PE150 reads at the HudsonAlpha Institute for Biotechnology in Huntsville, AL USA.

**Table 1:** Overview of PacBio HiFi and Omni-C sequencing data generated for the ‘Honeycrisp’ genome assembly.

| Library | Sequencing | Length (Nucleotides) | Number of reads |
| --- | --- | --- | --- |
| JNQN | Omni-C | 150 | 951,241,272 |
| HiFi-1 | PacBio HiFi | 14,881 * | 1,088,992 |
| HiFi-2 | PacBio HiFi | 14,429 * | 1,454,526 |

\* Average length

**Table 2:** Summary of ‘Honeycrisp’ genome assembly statistics.

| Assembly | Length | # Contigs | Longest contig | N50 | L50 | QV | k-mer completeness (%) | BUSCO (%) |
| --- | --- | --- | --- | --- | --- | --- | --- | --- |
| Honeycrisp Haplome A | 674,476,353 | 473 | 55,653,390 | 32,818,622 | 9 | 64.5 | 82.7 | 98.6 |
| Honeycrisp Haplome B | 660,238,068 | 215 | 56,154,892 | 31,578,807 | 9 | 66.7 | 83 | 98.7 |
| Combined |  |  |  |  |  | 65.5 | 98.6 |  |

**Table 3:** Yield of Illumina transcriptome sequencing of fruit, leaves, and flower tissues of apples and pear generated and used for genome annotation in this study.

| Cultivar | Tissue | Reads | Yield (Gb) | Yield P20 (Gb) | Ave Read length | NCBI SRA |
| --- | --- | --- | --- | --- | --- | --- |
| Honeycrisp | Fruitlet stage 1 | 45,773,784 | 13,823,682,768 | 13,069,822,280 | 142 | SAMN29611971 |
|  | Fruitlet stage 2 | 35,618,706 | 10,756,849,212 | 10,227,275,771 | 143 | SAMN29611972 |
|  | Budding leaves | 81,448,971 | 24,597,589,242 | 22,769,634,770 | 139 | SAMN29611973 |
|  | Expanding leaves | 35,381,039 | 10,685,073,778 | 9,971,308,535 | 141 | SAMN29611974 |
|  | Half-inch terminal buds | 47,811,924 | 14,439,201,048 | 13,409,542,519 | 140 | SAMN29611975 |
|  | Flower buds | 45,822,773 | 13,838,477,446 | 13,175,876,315 | 144 | SAMN29611976 |
|  | Open flowers | 30,938,395 | 9,343,395,290 | 8,718,474,885 | 141 | SAMN29611977 |
| Gala | Fruitlet stage 1 | 80,440,219 | 24,292,946,138 | 22,928,129,883 | 142 | SAMN29611954 |
|  | Fruitlet stage 2 | 32,475,136 | 9,807,491,072 | 9,284,944,973 | 143 | SAMN29611955 |
|  | Budding leaves | 30,368,057 | 9,171,153,214 | 8,508,033,713 | 140 | SAMN29611956 |
|  | Expanding leaves | 40,650,277 | 12,276,383,654 | 11,306,267,120 | 138 | SAMN29611957 |
|  | Roots from tissue culture | 35,324,786 | 10,668,085,372 | 9,940,132,737 | 140 | SAMN29611958 |
|  | Quarter-inch terminal buds | 37,532,631 | 11,334,854,562 | 10,634,379,784 | 141 | SAMN29611959 |
|  | Flower buds | 39,636,821 | 11,970,319,942 | 11,141,652,382 | 140 | SAMN29611960 |
|  | Open flowers | 34,363,075 | 10,377,648,650 | 9,775,838,818 | 142 | SAMN29611961 |
| Red Delicious | Fruitlet stage 2 | 27,319,955 | 8,250,626,410 | 7,682,200,349 | 140 | SAMN29611962 |
| Granny Smith | Fruitlet stage 1 | 29,426,606 | 8,886,835,012 | 8,335,731,187 | 141 | SAMN29611963 |
|  | Fruitlet stage 2 | 72,205,133 | 21,805,950,166 | 20,663,261,900 | 143 | SAMN29611964 |
|  | Budding leaves | 57,244,195 | 17,287,746,890 | 16,179,280,911 | 141 | SAMN29611965 |
|  | Expanding leaves | 40,798,422 | 12,321,123,444 | 11,499,303,808 | 140 | SAMN29611966 |
|  | Roots from tissue culture | 32,493,822 | 9,813,134,244 | 9,207,784,729 | 141 | SAMN29611967 |
|  | Quarter-inch terminal buds | 30,394,263 | 9,179,067,426 | 8,512,945,196 | 140 | SAMN29611968 |
|  | Flower buds | 29,735,514 | 8,980,125,228 | 8,364,532,017 | 140 | SAMN29611969 |
|  | Open flowers | 34,303,317 | 10,359,601,734 | 9,603,420,430 | 140 | SAMN29611970 |
| WA 38 | Fruitlet stage 1 | 45,284,208 | 13,675,830,816 | 12,831,991,620 | 141 | SAMN29611978 |
|  | Fruitlet stage 2 | 25,486,256 | 7,696,849,312 | 7,261,195,330 | 142 | SAMN29611979 |
|  | Budding leaves | 39,339,589 | 11,880,555,878 | 11,017,185,994 | 140 | SAMN29611980 |
|  | Expanding leaves | 34,784,980 | 10,505,063,960 | 9,719,694,010 | 139 | SAMN29611981 |
|  | Roots from tissue culture | 33,935,508 | 10,248,523,416 | 9,426,506,860 | 138 | SAMN29611982 |
|  | Quarter-inch terminal buds | 88,677,165 | 26,780,503,830 | 24,913,194,030 | 140 | SAMN29611983 |
|  | Flower buds | 23,170,354 | 6,997,446,908 | 6,588,921,074 | 142 | SAMN29611984 |
|  | Open flowers | 35,274,250 | 10,652,823,500 | 9,941,466,644 | 141 | SAMN29611985 |
| d'Anjou | Fruitlet stage 1 | 89,462,306 | 27,017,616,412 | 25,459,693,894 | 142 | SAMN29611986 |
|  | Fruitlet stage 2 | 48,481,031 | 14,641,271,362 | 13,921,844,851 | 143 | SAMN29611987 |
|  | Budding leaves | 29,823,484 | 9,006,692,168 | 8,442,259,663 | 141 | SAMN29611988 |
|  | Expanding leaves | 57,920,009 | 17,491,842,718 | 16,460,531,509 | 142 | SAMN29611989 |
|  | Quarter-inch terminal buds | 40,966,825 | 12,371,981,150 | 11,476,090,088 | 140 | SAMN29611990 |
|  | Flower buds | 29,183,231 | 8,813,335,762 | 8,264,473,671 | 141 | SAMN29611991 |
|  | Open flowers | 32,128,369 | 9,702,767,438 | 8,996,878,963 | 140 | SAMN29611992 |

### Repeat analysis and gene annotation

Repetitive elements on both haplotypes were annotated using EDTA v2.0.0 [43] with flags “--genome, --anno 1, --sensitive=1”. To supplement *ab initio* gene predictions, extensive extrinsic gene annotation homology evidence is needed. Thus, we downloaded existing RNA-seq data for ‘Honeycrisp’ apples from NCBI using SRA toolkit v2.9.6-1 (SRX3408575, SRX5369275, SRX5369276, SRX5369290, SRX5369299, SRX5369300, SRX5369302, SRX8712695 and SRX8712718) [44–46], and combined with the RNA-seq data generated for this project (described above). We *de novo* assembled these two sets of RNA transcripts separately using Trinity v2.13.2 [47], where we used the flag --trimmomatic to filter the reads for quality. Because the newly generated RNA-seq data were strand-specific, for these we also used the flag “--SS_lib_type RF”. We identified open reading frames using TransDecoder v5.5.0 [48]. Gene annotation was performed using BRAKER2 v2.1.6 [49], where we ran BRAKER2 twice, with RNA-seq data and protein databases run separately. For the RNA-seq run, we first filtered the data for adapters and quality using TRIMMOMATIC v0.39 [50] with leading and trailing values of 3, sliding window of 30, jump of 10, and a minimum remaining read length of 40. We next mapped these data to the genome using STAR v2.7.9a [51] and combined the BAM files using SAMtools [52]. For the homology-based annotation in BRAKER2, we used gene models from *Malus domestica* ‘Gala’ diploid v2, *M. sieversii* diploid v2 [27], *M. baccata* v1 [53]. *M. domestica* ‘Golden Delicious’ double haploid v1 (GDDH13) [29], *Pyrus communis* ‘Barlett’ double haploid v2 [54], and our *de novo* assemblies, in addition to the viridiplantae OrthoDB [55]. We filtered the resulting AUGUSTUS [49] output for those that contained full hints (gene model support) and combined the two runs using TSEBRA v1.0.3 [56]. Finally, we removed any transcript/gene that had ≥90% softmasking, *i.e.,* mainly repeat sequences. Genome annotation completeness of our genome and other *Malus* genomes were assessed using BUSCO v5.2.2 [40] with the “eudicots_odb10” database for comparative purposes.

The final ‘Honeycrisp’ gene sets from both haplomes were annotated with InterProScan v5.44-79.0 [57,58], including a search against all the available interpro databases and Gene Ontology (GO) [59,60] prediction. In addition, genes were searched against the 26Gv2.0 OrthoFinder v1.1.5 [61] gene family database using both BLASTp [62] and HMMscan [63] classification methods with the *GeneFamilyClassifier* tool from PlantTribes 2 (github.com/dePamphilis/PlantTribes/). This analysis provided additional functional annotation information that includes gene counts of scaffold taxa, superclusters at multiple clustering stringencies, and functional annotations that were pulled from various public genomic databases.

### Comparative genomics

Similarities in lengths and structural variations between the two haplomes were determined by running MUMmer v4.0 [64] and Assemblytics [65]. To identify the shared and unique gene families among *Malus* species and cultivars, genes from the six publicly available *Malus* genomes (Table 5) were integrated into the aforementioned PlantTribes 2 gene model database (26Gv2.0) using the same method as described above. The overlapping orthogroups (with at least 30 counts in the category) among the eight *Malus* annotations (including both haplomes from ‘Honeycrisp’) were calculated and visualized with an upset plot generated by TBtools v1.0986982 [66].

**Table 4:** Summary of repetitive element annotation in Haplome A and Haplome B of the ‘Honeycrisp’ genome assemblies.

| Class |  | Haplome A | Haplome B |
| --- | --- | --- | --- |
| LTR |  |  |  |
|  | Copia | 9.73% | 9.60% |
|  | Ty3 | 20.29% | 17.80% |
|  | unknown | 14.89% | 16.86% |
| TIR |  |  |  |
|  | CACTA | 2.21% | 1.95% |
|  | Mutator | 4.16% | 4.25% |
|  | PIF Harbinger | 2.43% | 2.60% |
|  | Tc1_Mariner | 0.15% | 0.27% |
|  | hAT | 2.30% | 2.31% |
|  | polinton | -- | 0.01% |
| nonLTR |  |  |  |
|  | LINE_element | 0.18% | 0.17% |
|  | unknown | 0.09% | 0.18% |
| nonTIR |  |  |  |
|  | helitron | 2.95% | 3.18% |
| repeat region |  | 2.91% | 2.78% |
| **Total** |  | **62.43%** | **61.97%** |

**Table 5:** Comparison of genomic features and assembly statistics of current assembly of ‘Honeycrisp’ genome and previously published genomes of apples.

| Genomes | ‘Honeycrisp’ | ‘Gala’,<br>*M. sieversii*,<br>*M. sylvestris*<br>(all Diploid) | HFTH1;<br>‘Hanfu’<br>(Triploid) | GDDH13;<br>‘Golden Delicious’<br>(Double haploid) | ‘Golden<br>Delicious’<br>(Diploid) |
| --- | --- | --- | --- | --- | --- |
| Reference | This work | Sun et al. 2021 | Zhang et. 2019 | Daccord et al. 2017 | Velasco et al. 2010 |
| Assembly |  |  |  |  |  |
| Haploid genome size (Mb) | 660-674 | 666-679 | 658.9 | 651 | 742 |
| scaffold N50 | 31.6-32.8 | 6.1-21.8 | 6.99 | 5.5 | 16Kb |
| Complete BUSCO | 98.6-98.7% | 98.0-98.8% | 98.6% | 98.0% | 82.0% |
| **Annotation** |  |  |  |  |  |
| **Protein-coding genes** | 47,563-48,655 | 44,691-44,847 | 44,677 | 42,140 | 57,386 |
| **Complete BUSCO** | 96.8-97.4% | 94.6-95.4% | 93.6% | 96.1% | 68% |
| **Gene family** |  |  |  |  |  |
| **Number of OG in 26Gv2** | 10,351-10,367 | 10,044-10,115 | 9,974 | 10,117 | 8,824 |

## Results

### A haplotype-phased chromosome-scale assembly

In total, nearly 55X coverage of PacBio HiFi reads and nearly 200X coverage of Dovetail Omni-C reads (Table 1) was generated. This included 2,543,518 HiFi reads with an average length of 14,655 bp and ~91% of reads ≥10,000 bp. Two phased haplomes, haplome A (HAP1) and haplome B (HAP2, these two sets of terms will be used interchangeably in this manuscript), were assembled and validated by inspection of the Omni-C contact maps (Figure 2). Both haplomes are highly contiguous and of similar size. HAP1 is 674 Mb in length, contained in 473 contigs with a contig N_50_ of 32.8 Mb, whereas HAP2 is 660 Mb in length, contained in 215 contigs with a contig N_50_ of 31.6 Mb (Table 2). Zero miss-joins requiring manual breaks were identified in the assemblies. For HAP1, a total of 13 joins were made to build the final assembly into 17 chromosomes, with 95.4% of the assembled sequence contained in the 17 pseudomolecules representing chromosomes. A total of 19 joins were made for HAP2, with 98.2% of the assembled sequence in the 17 pseudomolecules. Based on the Merqury k-mer analysis (Figure 3), the HAP1 assembly had a k-mer completeness of 82.7% (Quality value (QV) 64.5), the HAP2 assembly 83% (QV 66.7), and the combined assemblies were 98.6% (QV 65.5) (Table 2). BUSCO completeness of HAP1 was 98.6% and HAP2 98.7%, suggesting high genome completeness for both haplomes, comparable or superior to other high quality apple genome assemblies (Table 5). The two haplomes are structurally similar to each other (Figure 4). Compared to the assembly statistics of previously published apple genomes, the current ‘Honeycrisp’ assemblies are the most contiguous to date (Table 5).

**Figure 2:**
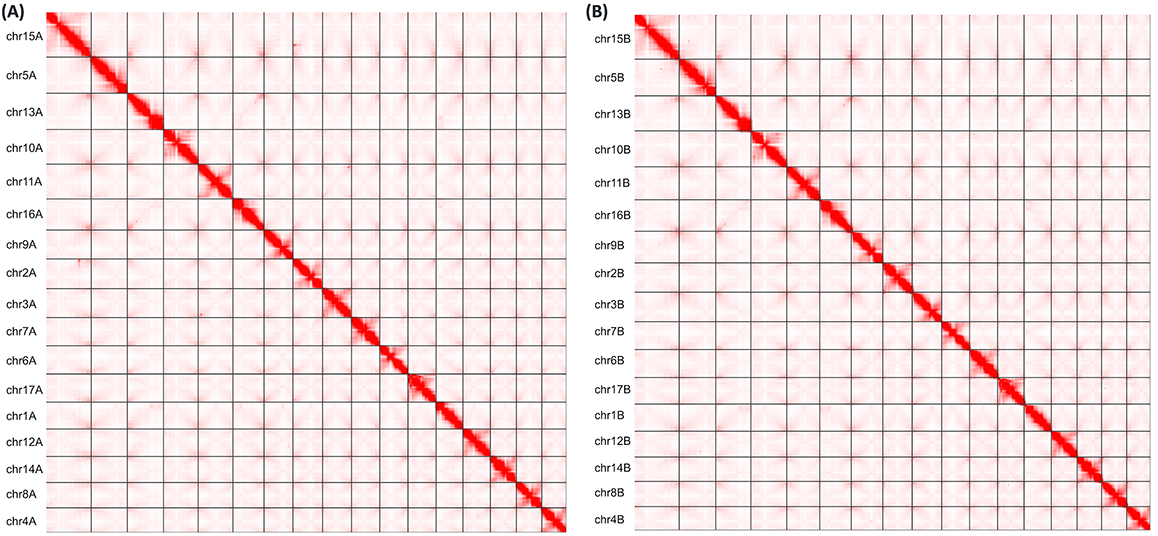
Omni-C contact maps of the assembled chromosome-length scaffolds of 17 chromosomes of (A) Haplome A and (B) Haplome B, from the ‘Honeycrisp’ genome.

**Figure 3:**
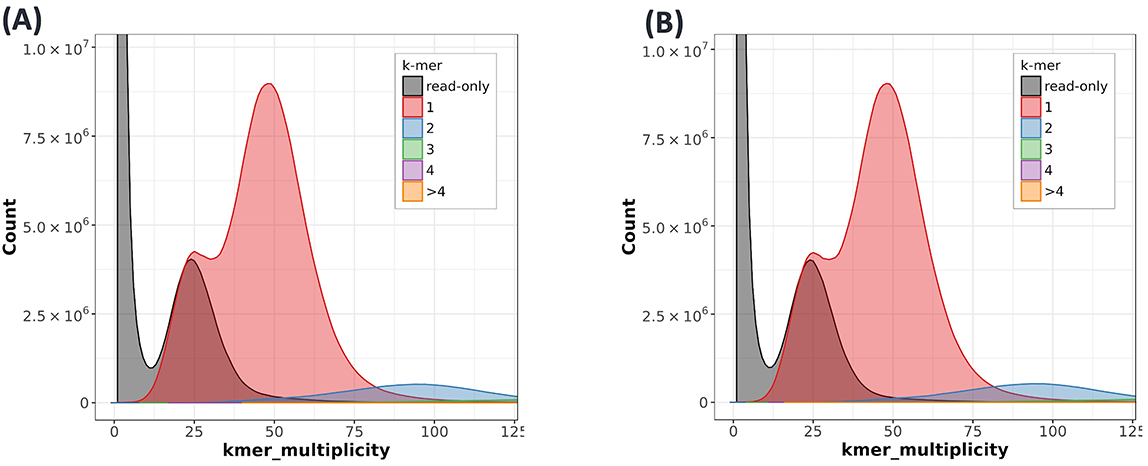
Histogram of k-mer multiplicity of sequence reads for (A) Haplome A and, (B) Haplome B of ‘Honeycrisp’ genome assemblies. K-mer multiplicity (x-axis) is plotted against k-mer counts (y-axis) to estimate the heterozygosity, copy numbers, sequencing depth, and completeness of a genome using Merqury v1.3 [41]. Colors in the plot represent the number of times each k-mer is found in the genome assembly.

**Figure 4:**
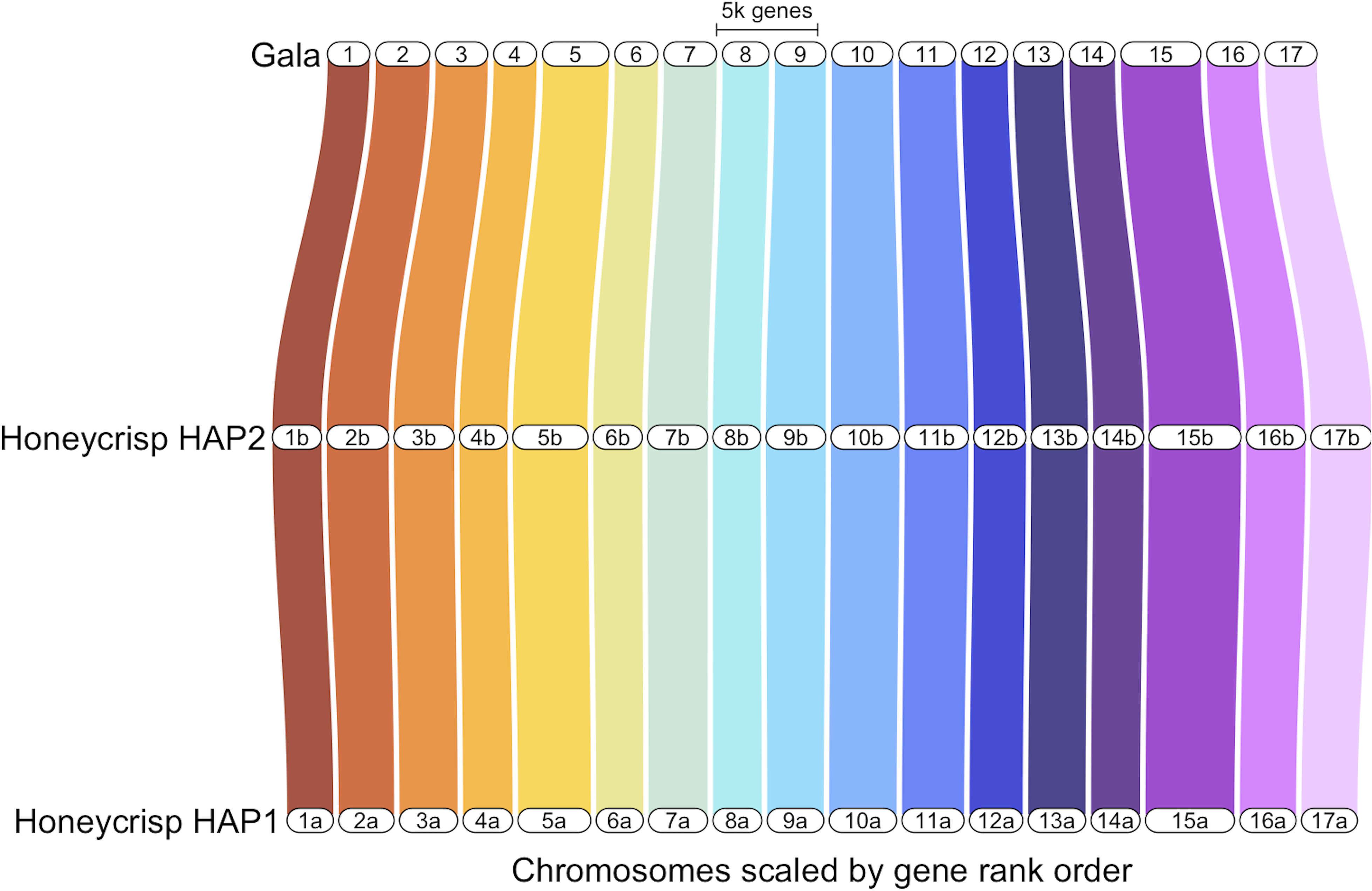
Synteny comparison of ‘Honeycrisp’ Haplome 1 (HAP1), ‘Honeycrisp’ Haplome 2 (HAP2) from this study, and ‘Gala’ [27] genomes. GENESPACE [39] was used for synteny comparison.

### Genome annotation

The yield of Illumina transcriptome sequencing data of fruit, leaves, and flower tissues of apples and pear ranged from approximately 9 to 27 gigabases (Gb) in flowers and leaf buds respectively (Table 3). Nearly 62% of both haplomes were annotated as repetitive DNA, mostly comprised of Long Terminal Repeat (LTR) retrotransposons (Table 4). A total of 47,563 genes were annotated in HAP1 and 48,655 in HAP2, slightly more than in other published *Malus* annotations (Table 5). Complete BUSCO scores of the protein annotations are 96.8% for HAP1 and 97.4% for HAP2, the highest completeness among all publicly available *Malus* genome annotations (Table 5). 72.85% and 68.88% of the predicted transcripts were annotated with Interpro terms, 68.58% and 64.94% with Pfam domains, and 51.04% and 48.76% with at least one GO terms in HAP1 and HAP2, respectively. In the PlantTribes 2 classification, 91.11% and 85.50% of the predicted transcripts from HAP1 and HAP2, respectively, were assigned to pre-computed orthogroups.

As the number of plant genomes are being generated at an unprecedented speed, we developed the following gene naming convention to avoid potential ambiguity. Maldo.hc.v1a1.ch10A.g00001.t1: Maldo for *Malus domestica*; hc for the cultivar, ‘Honeycrisp’; v1a1 indicating this is the first assembly and first annotation of this genome; ch10A identifies that the gene is annotated from chromosome 10 (versus from an unplaced scaffold, which will be indicated by “sc”) in haplome A (HAP1) (versus haplome B (HAP2)); g00001 is a five digit gene identifier; t1 represents a transcript number of the gene.

### Gene family analysis

A gene family evaluation was performed using PlantTribes 2 and its 26Gv2-scaffold orthogroup database, which contains representative protein coding sequences from most major land plant lineages. A total of 11,263 unique orthogroups (OGs) were identified in all eight *Malus* annotations (including the two ‘Honeycrisp’ haplomes) investigated. ‘Honeycrisp’ transcripts were assigned to 10,351 and 10,367 orthogroups, similar to ‘Gala’ and GDDH13 (Table 5 and Figure 5). We further investigated orthogroups that are shared and unique in the eight *Malus* annotations. A vast majority (7,645) of orthogroups are shared by all the genomes, and a total of 9,279 orthogroups were shared among both ‘Honeycrisp’ haplomes and five other genomes (Figure 5). This comparison indicates that the ‘Honeycrisp’ annotation captured genes in virtually all the *Malus* gene families. In addition, we also found 54 orthogroups that are unique to ‘Honeycrisp’ (*i.e.,* shared by the two ‘Honeycrisp’ haplomes only) and 35 and 32 that are unique to each ‘Honeycrisp’ haplome (Figure 5). These orthogroups could provide valuable information in the molecular mechanisms underlying genotype-specific traits.

**Figure 5:**
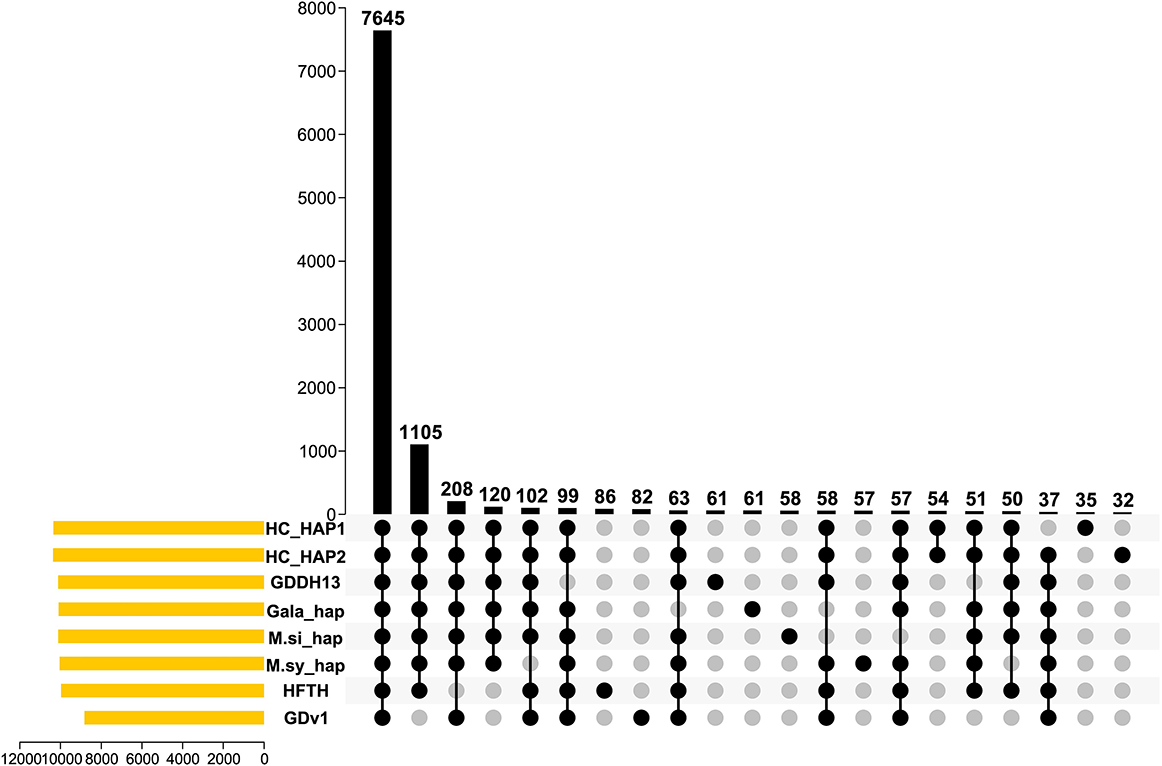
The Honeycrisp genome captured a vast majority of *Malus* gene families. Black dots indicate presence of gene families and gray dots indicate absence. Yellow horizontal bars represent the number of orthogroups in each genome. The black vertical bars represent the number of orthogroups in each category. Genome abbreviations-HC: ‘Honeycrisp’ (this work); GDDH13: *Malus domestica* GDDH13; Gala_hap: *M. domestica* ‘Gala’ haploid; M.si_hap: *M. sieversii* haploid; M.sy_hap: *M. sylvestris* haploid; HFTH: *M. domestica* HFTH1; GDv1: *M. domestica* Golden Delicious v1.

## Re-use Potential

This fully phased, high-quality, chromosome-scale genome of ‘Honeycrisp’ apple will add to the toolbox for apple genetic research and breeding. It will enable genetic mapping, identification of genes, and development of molecular markers linked to disease, pest resistance, abiotic stress tolerance and adaptation, as well as horticulturally relevant harvest and postharvest fruit quality traits for use in apple breeding programs. Ultimately, the addition of high-quality genomic resources for ‘Honeycrisp’ can lead to enhanced orchard and supply chain management for many other apple cultivars, promoting future sustainability of the pome fruit industry.

## Data Availability

The whole genome sequence data generated in this study have been deposited at the National Center for Biotechnology Information (NCBI) database under BioProject ID PRJNA791346. PacBio HiFi reads, and Hi-C reads are deposited in NCBI with the SRA accession number SAMN24287034 and SAMN29611953, respectively. Transcriptomic data generated in this study for genome annotation are deposited in NCBI with SRA accession number from SAMN29611954 to SAMN29611992. The Maldo.hc.v1a1 ‘Honeycrisp’ genome assembly, gene annotation, and functional annotation for both haplomes can be accessed *via* the Genomic Database for Rosaceae (in progress) and the GigaScience GigaDB repository.

## Declarations

### List of abbreviations

BLAST: Basic Local Alignment Search Tool; bp: Base Pair; BUSCO: Benchmarking Universal Single Copy Orthologs; Gb: gigabases; GO: Gene Ontology; HMW: High-molecular-weight; JBAT: Juicebox Assembly Tools; LTR: Long Terminal Repeat; NCBI: National Centre for Biotechnology Information; OG: Orthogroup; QV: Quality Value; RIN: RNA Integrity Number; SMRT: Single Molecule Real Time; TRF: Tandem Repeat Finder

### Author Contribution

A.K., and L.H. conceptualized, designed, and managed the project. S.B.C., H.Z., A.H., and H.H, constructed DNA and RNA, and RNA-seq libraries for sequencing. S.B.C., and H.Z., performed genome and transcriptome sequence analysis and interpretation. S.B.C., H.Z., A.S., H.H., A.H., L.H., and A.K. drafted, revised, and finalized the manuscript. All authors read and approved the manuscript.

## Acknowledgements

We acknowledge the help of Della Cobb-Smith and Jugpreet Singh for sample collection and preparation for genomic DNA extraction.

